# The seagrass methylome memorizes heat stress and is associated with variation in stress performance among clonal shoots

**DOI:** 10.1101/787754

**Authors:** A Jueterbock, C Boström, James A Coyer, JL Olsen, M Kopp, AKS Dhanasiri, I Smolina, S Arnaud-Haond, Y Van de Peer, G Hoarau

**Affiliations:** Marine Molecular Ecology Group, Faculty of Biosciences and Aquaculture, Nord University, Universitetsalleen 11, 8026 Bodø, Norway; Environmental and Marine Biology, Åbo Akademi University, Artillerigatan 6 FI-20520 Åbo, Finland; Shoals Marine Laboratory, University of New Hampshire, Durham, NH 03824, USA; Ecological Genetics-Genomics Group, Groningen Institute of Evolutionary Life Sciences, University of Groningen, 9747 AG Groningen, The Netherlands; UMR MARBEC, Université de Montpellier, Ifremer, IRD, CNRS, Avenue Jean Monnet, CS 30171, 34203 Sète Cedex, France; Department of Plant Biotechnology and Bioinformatics, Ghent University and VIB Center for Plant Systems Biology, Technologiepark 71, 9052, Ghent, Belgium; Center for Microbial Ecology and Genomics, Department of Biochemistry, Genetics and Microbiology, University of Pretoria, Pretoria, South Africa

**Keywords:** DNA methylation, ecological epigenetics, clonality, heat stress, seagrass, *Zostera marina* (eelgrass), priming

## Abstract

Evolutionary theory predicts that clonal organisms are more susceptible to extinction than sexually reproducing organisms, due to low genetic variation and slow rates of evolution. In agreement, conservation management considers genetic variation as the ultimate measure of a population’s ability to survive over time. However, clonal plants are among the oldest living organisms on our planet. Here, we test the hypothesis that clonal seagrass meadows display epigenetic variation that complements genetic variation as a source of phenotypic variation. In a clonal meadow of the seagrass *Zostera marina* we characterized DNA methylation among 42 shoots. We also sequenced the whole genome of 10 shoots to correlate methylation patterns with photosynthetic performance under exposure to, and recovery from 27°C, while controlling for somatic mutations. Here, we show for the first time that clonal seagrass shoots display DNA methylation variation that is associated with variation in fitness-related traits: photosynthetic performance and heat stress resilience. The co-variation in DNA methylation and phenotype may be linked via gene expression because methylation patterns varied in functionally relevant genes involved in photosynthesis, and in the repair and prevention of heat-induced protein damage. A >five week epigenetic heat stress memory may heat-harden previously heat-exposed shoots. While genotypic diversity has been shown to enhance stress resilience in seagrass meadows, we suggest that epigenetic variation plays a similar role in meadows dominated by a single genotype. Consequently, conservation management of clonal plants should consider epigenetic variation as indicator of resilience and stability, and restoration efforts may benefit from stress-priming transplanted seeds or shoots.

## 1 Introduction

Genetic variation is considered key to long-term survival of populations (Bijlsma and Loeschcke, 2012), and is recognized as a main target for the conservation and restoration of biological diversity (Lande, 1988; Spielman et al., 2004; Laikre et al., 2009). In contrast, lack of genetic variation is regarded as an evolutionary dead-end (Lynch and Lande, 1998) but clonal growth challenges the expected relationship between genetic diversity and long-term survival. For example, conservation management follows the rough guideline that within a population >1,000 genetically different individuals must mate randomly to avoid inbreeding depression, and retain evolutionary potential (Frankham et al., 2014). Yet, roughly 40% of all plants can reproduce asexually (Tiffney and Niklas, 1985), mostly by clonal growth where parental genotypes (genets) grow vegetative modules (ramets) that often remain connected via underground stolons or rhizomes.

Although clones benefit from resource sharing, niche specialization, and rapid vegetative growth (Liu et al., 2016), they are predicted to survive only for short periods, and in stable environments (Silvertown, 2008). Asexual reproduction is assumed to lead to slow rates of genetic evolution, and the lack of DNA repair mechanisms afforded by meiosis (Muller’s ratchet, mutational meltdown) (Muller, 1964; Gabriel et al., 1993; Lynch et al., 1993). Despite asexual reproduction, many of our most important crops are clones (McKey et al., 2010), including banana, garlic, hops, potatoes, and turmeric, as well as many of the earth’s most invasive and oldest plants. For example, genets of Palmer’s oak (*Quercus palmeri*) and seagrass (*Posidonia oceanica*) are estimated older than 10,000 years (May et al., 2009; Arnaud-Haond et al., 2012) Thus, long-term survival appears not to rely solely on sexual production.

Somatic mutations can create a certain level of genetic diversity, and may explain some evolutionary potential of clonal organisms (Whitham and Slobodchikoff, 1981; Loxdale et al., 2003; Lushai et al., 2003; Reusch and Boström, 2011). For example, ~7,000 single nucleotide polymorphisms (SNPs), 597 in coding regions and 432 non-synonymous, distinguish ramets of a large Finnish clone of the seagrass *Zostera marina* (Yu et al., 2020). To set this in perspective,139,321 biallelic SNPs were reported in coding regions among four populations of the same species (Jueterbock et al., 2016). The degree to which epigenetic variation can contribute to phenotypic heterogeneity in ecologically relevant traits, independently from the underlying genetic variation, is a key question in assessing its contribution to stress tolerance, and long-term survival of clonal organisms.

The definition of epigenetics is currently heavily debated (Ptashne, 2007; Greally, 2018). Here, ‘epigenetic’ implies molecular variations that do not alter the DNA sequence but have the potential to change gene expression, and include non-coding RNAs (ncRNAs), histone modifications, and DNA methylation (Bossdorf et al., 2008). DNA methylation is, from an evolutionary perspective, the most relevant epigenetic mechanism because it can be independent from genetic variation (Bossdorf et al., 2008; Schmitz et al., 2013; Kilvitis et al., 2014), and transgenerationally stable (Boyko et al., 2010; Verhoeven et al., 2010; Ou et al., 2012; Bilichak et al., 2015; Williams and Gehring, 2017). DNA methylation involves the addition of a methyl-group to the C5 position of a cytosine in DNA sequence motifs (CG, CHG, and CHH in plants, where H stands for A, C, or T) (Kilvitis et al., 2014). Depending on sequence context, methylation can be associated with gene activation or silencing (Bossdorf et al., 2008; Niederhuth and Schmitz, 2017). While CG methylation in gene bodies often correlates with increased gene expression, methylation in promoters and repeat regions, such as transposable elements (TEs), silences expression (Feng et al., 2010; Seymour et al., 2014; Dubin et al., 2015; Bewick and Schmitz, 2017; Zhang et al., 2018).

The methylome, or set of DNA methylation modifications in an organism’s genome, can change spontaneously at a rate of 2.5×10^−4^ to 6.3×10^−4^ methylation polymorphisms per CG site per generation, which is about 7×10^4^ higher than the genetic mutation rate of base substitutions per site per generation (Schmitz et al., 2011; van der Graaf et al., 2015). That methylome variation can enhance productivity, and pathogen resistance, has been shown in *Arabidopsis thaliana* plant populations (Latzel et al., 2013). Moreover, methylation variation explained the rapid invasive success of the Japanese knotweed (*Fallopia japonica*) by facilitating differentiation in response to new habitats despite decreased genetic variation (Richards et al., 2012). This suggests that methylation variation complements genetic variation as a source of phenotypic variation in plant populations deprived of genotypic diversity (Gao et al., 2010; Richards et al., 2012; Verhoeven and van Gurp, 2012; Latzel et al., 2013; Zhang et al., 2013; Vanden Broeck et al., 2018).

Unlike genetic variants, methylation variants can also switch state directly in response to the environment (Dowen et al., 2012) and, if stable enough, establish a molecular memory that can be involved in stress priming. Priming, also referred to as hardening or conditioning, is well studied in terrestrial plants (Hilker et al., 2016; Lämke and Bäurle, 2017; Pawar and Laware, 2018), and refers to the plants’ ability to acquire a stress memory that enhances performance under second stress exposure by responding faster, stronger, or in response to a lower threshold as compared with naïve plants (Balmer et al., 2015; Lämke and Bäurle, 2017). Stress responsive methylation marks that are transgerationally stable can create adaptive phenotypic changes within a single generation (Bossdorf et al., 2008; Jablonka and Raz, 2009; Nicotra et al., 2010; Verhoeven et al., 2010; Hirsch et al., 2012; Richards et al., 2012; Douhovnikoff and Dodd, 2014; Verhoeven and Preite, 2014).This may explain why in some cases a primed state can improve plant performance even across generations (Rasmann et al., 2012; Slaughter et al., 2012; Kuźnicki et al., 2019).

Under clonal growth, DNA methylation patterns are expected to be more stably inherited than under sexual reproduction (Verhoeven and Preite, 2014), because clonal growth circumvents epigenetic reprogramming during gameto- and embryogenesis. Although stable transmission across asexual generations has been shown for environment-specific phenotypes and DNA methylation patterns in clonal plants (Verhoeven et al., 2010, 2018; Verhoeven and van Gurp, 2012; Vanden Broeck et al., 2018), no link has been demonstrated between methylation variation and fitness-related traits. Thus, while epigenetic mechanisms have been suggested to contribute to clonal plant success (Douhovnikoff and Dodd, 2014; Verhoeven and Preite, 2014; Dodd and Douhovnikoff, 2016; Latzel et al., 2016), empirical evidence is virtually lacking.

Clonal propagation is especially well-developed in aquatic plants (Barrett, 2015). Seagrasses, the only plants to inhabit the marine world, form the foundational basis of some of the most productive and highly diverse coastal marine ecosystems on the planet, and are essential for the health and abundance of economically exploited marine species (Costanza et al., 1997; Larkum et al., 2006; Orth et al., 2006). Ecosystem services are worth more than € 16,000 ha^−1^ year^−1^ (Costanza et al., 1997), including nursery grounds, habitat and food for fish and invertebrates, protection of the coastline from erosion, carbon sequestration of up to 186 g C m^2^ yr^−1^ (Fourqurean et al., 2012; Duarte et al., 2013), and nutrient fixation (Orth et al., 2006; Procaccini et al., 2007).

Over the last decades, losses of seagrass ecosystems have been documented worldwide due to increasing anthropogenic stressors such as invasive species, sediment and nutrient runoff, dredging, aquaculture, rising sea levels, and global warming (Orth et al., 2006; Chefaoui et al., 2018). Losses are expected to accelerate under projected global temperature increase, as ocean warming is considered the most severe threat among climate change factors (Repolho et al., 2017; Duarte et al., 2018) and seagrass meadows have tracked temperature changes in the past (Olsen et al., 2004). How accurate these shifts and losses can be predicted depends on our knowledge of drivers of adaptive potential, including epigenetic diversity (Duarte et al., 2018).

In this study, we characterize the functional relevance of epigenetic variation for heat stress resilience in the seagrass *Zostera marina*. *Z*. *marina* is the most widely distributed seagrass in the northern hemisphere, inhabiting highly contrasting habitats from sub-arctic to sub-tropical waters (Green and Short, 2004; Olsen et al., 2004; Boström et al., 2014). Few plants display such dramatic range in clonal diversity as *Z*. *marina* (Reusch et al., 2000; Olsen et al., 2004). Its clonal architecture varies from genetically diverse meadows with high levels of sexual reproduction, to meadows composed of a single large clone due to exclusive vegetative reproduction (Reusch et al., 1999; Olsen et al., 2004; Becheler et al., 2010; Reusch and Boström, 2011). Clonality of *Z*. *marina* peaks in the Baltic Sea, where large clones were estimated > 1,000 years old (Reusch et al., 1999). These meadows display remarkable phenotypic variability and persistence across space and time, under perturbations (e.g. high temperatures) that represent environmental stresspredicted elsewhere for the future (Reusch et al., 2018). Thus, clonal meadows (such predominated by a single genet) confound experimental results showing a positive effect of genotypic (i. e. clonal) richness on the productivity and stress resilience of *Z*. *marina* (Hughes and Stachowicz, 2004; Reusch et al., 2005; Reusch and Hughes, 2006; Ehlers et al., 2008; Hughes et al., 2008). In other words, if stress resilience and tolerance would rely strongly on genotypic diversity, clones would not be able to survive for such long time periods.

The resilience, longevity, and adaptive potential of clonal seagrass meadows remain unknown without a fundamental analysis of their epigenetic variation and its ecological relevance. Therefore, this study tests the hypothesis that variation in DNA methylation can promote functional phenotypic diversity and, thus, may explain how clonal seagrass meadows can persist over millennia.

Specific objectives were to: 1) Characterize DNA methylation variation in an ancient clonal meadow of *Z*. *marina* (>1,000-years-old); 2) Identify the functional role of this variation in heat-stress performance–specifically how photosynthetic performance is linked to DNA methylation variation independently from underlying somatic mutations, and 3) Assess to what extent heat-responsive methylation patterns are reset versus memorized after the stress is removed.

## 2 Materials and Methods

### 2.1 Sampling and cultivation

We sampled seagrass shoots from a clonal meadow (Reusch et al., 1999) in the Baltic Sea (Åland Islands, 60°09’50.4”N, 19°31’48.1”E) in 2015 (Figure 1) by collecting 2-3 shoots attached to the same rhizome (ramets) every 3 meters along a 250 m transect (Supplementary table S1). Shoots were transported in seawater-filled cooling boxes to the field station at Nord University, Norway. Leaf tissue of 42 shoots, randomly chosen from the 2-3 connected ramets, was flash frozen in liquid nitrogen for subsequent DNA extraction (Supplementary table S1).

**Figure 1.**
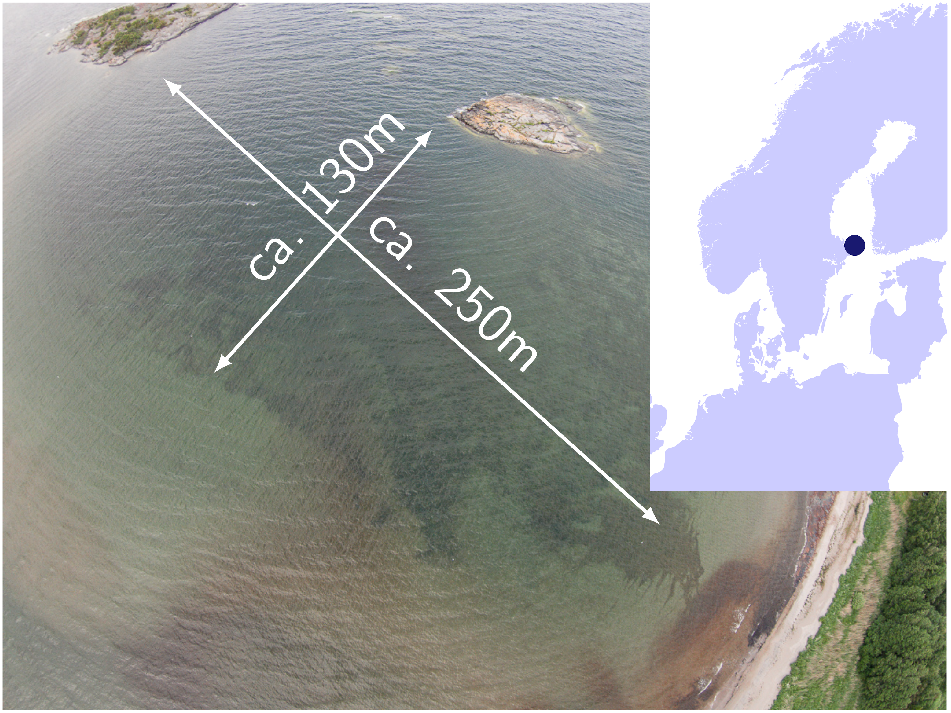
Baltic Sea sampling site of a > 1,000 years old clonal meadow where 42 *Zostera marina* shoots were collected along a 250 m transect.

Ten ramets of the same genet (Supplementary table S1) were replanted in plastic containers (10×15 cm) filled with sand from the sampling site, placed in a 1,280 L aquarium at 15°C, and illuminated with a 16:8 h light:dark cycle, and 200-220 μmol m^−2^ s^−1^ (OSRAM Fluora, 150 W). Seawater at 5.5 PSU, corresponding to the salinity at the collecting site, was obtained by mixing freshwater and natural filtered seawater at 32 PSU. The water, filled to a 40 cm level, was kept in constant motion with airstones. Once a week, 50% of the water was renewed after removing epiphytic algae.

### 2.2 Heat stress experiments

After two weeks of acclimation, the 10 clonal ramets were exposed to heat stress (Supplementary table S1) in climate chambers (Fitotron, weisstechnik), where they were distributed in aquaria (60 L) filled with aerated brackish water (5.5 PSU) to 30 cm, of which 50% was weekly renewed. Light was kept at 240 μmol m^−2^ s^−1^ and a 16:8 h light:dark cycle. The temperature in the climate chambers was increased with a daily increment of 3°C from 15°C to 27°C, which can be lethal for *Z*. *marina* (Greve et al., 2003), but is reached in the Baltic Sea during summer heat waves. After three weeks at 27°C, the temperature was decreased by a daily increment of 3°C from 27°C to 15°C, after which all shoots were returned to the 1,280 L aquarium at 15°C for a recovery period of 5.5 weeks.

At three time points (control, stress, recovery), one leaf of each shoot was excised and flash-frozen in liquid nitrogen. Control samples were collected at 15°C on the day before the temperature was increased. Stressed samples were collected at 27°C on the day before the temperature was decreased. Recovery samples were collected after the 5.5 week recovery period at 15°C.

### 2.3 Photosynthetic performance

At control, stress, and recovery time points, we measured for each shoot (two measurements for each of the two inner leaves) the increase in chlorophyll a fluorescence upon illumination after a 10 min dark period (OJIP curve) (Bussotti et al., 2010), with a PAM-Fluorometer (FluorPen FP100, Photon Systems Instruments) using a saturating pulse of 75% at 455 nm. From the measurements, we extracted the performance index PiABS (Strasser et al., 2000), reflecting the functionality of photosystem II (PSII) and photosynthetic performance in general (Živčák et al., 2008; Bussotti et al., 2010; Stefanov et al., 2011). We tested for correlation of PiABS values (mean values per shoot) between time points using a two-sided Pearson’s product moment correlation test in R v3.4.4 (R Core Team, 2019).

### 2.4 DNA extraction

All flash-frozen tissue was freeze-dried for one day, then stored at −80°C until DNA extraction. DNA was extracted with the HP Plant DNA mini kit (Omega Bio-Tek, protocol version May 2013) from ≥ 5 mg of lyophilized leaf tissue after grinding with a mixer-mill (Retsch MM 400) in 2 mL Eppendorf tubes with tungsten beads supplied (60 s at 30 Hz). We added 10 μL beta-mercaptoethanol at step 2, and equilibrated the columns at step 8 of the standard protocol. The extracted DNA was eluted in 2 x 100 μL EB buffer (Qiagen), cleaned and concentrated with the Clean and Concentrator-5 kit (Zymo Research, protocol v1.2.1) using 15,000 g for all centrifugation steps, and finally eluted in 2 x 30 μL EB Buffer (Qiagen) at 60°C.

### 2.5 Clonality

In order to identify ramets belonging to the same genet, we genotyped each shoot for seven microsatellite loci (ZosmarGA2, -GA3, -GA6, -CT3, -CT12, -CT19, -CT20) (Reusch and Boström, 2011). PCR was performed using a Veriti 96-Well Thermal Cycler (Applied Biosystems, Life Technologies) in two 10 μL multiplex reactions containing 2 μL cleaned genomic DNA, and 1X AccuStart II PCR ToughMix (Quanta bio). One multiplex reaction contained forward and reverse primers GA2, GA3, and GA6 at 0.5 μM each, and ran at 94°C for 4 min, followed by 30 cycles of 94°C for 60 s, 55°C for 90 s, and 72°C for 90 s, and a final extension at 72°C for 10 min. The other multiplex reaction contained forward and reverse primers CT3 (0.5 μM), CT12 (0.3 μM), CT19 (0.3 μM), and CT20 (0.3 μM), and ran at 94°C for 3 min, followed by 35 cycles of 94°C for 60 s, 57°C for 60 s, and 72°C for 60 s, and a final extension at 72°C for 10 min.

DNA fragment lengths were determined on an ABI 3500xl Genetic Analyzer from 1 μL of 1:99 diluted PCR products mixed with 8.9 μL of HiDi Formamide (Life Technologies), and 0.1 μL of Gene Scan 500 LIZ Size Standard (Life Technologies) after 5 min denaturation at 95°C. Alleles were called with the GeneMapper v4.1 Software (Applied Biosystems, Thermo Fisher Scientific). The shoots were assigned to multi-locus genotypes using the R package ‘RClone’ (Bailleul et al., 2016).

### 2.6 Whole genome sequencing, SNP detection, and genetic distance

In order to detect SNPs resulting from somatic mutations in the 10 heat-stressed ramets, genomic DNA libraries were prepared according to the TruSeq DNA PCR-Free (Illumina) protocol, and sequenced on one Illumina HiSeq 3/4000 lane (2×150bp) at the Norwegian Sequencing Centre (University of Oslo, Norway). Raw reads (25.7 Million to 44.3 Million per library, Supplementary Table S2, NCBI BioProject PRJNA575339^1^) were quality-checked with FastQC v0.11.8^2^ to control for aberrant read base content, length distribution, duplication, and over-representation. We used TrimGalore! v0.6.0^3^ to remove adapter sequences with a stringency of 3 bp overlap, and low-quality bases with a Phred score Q < 20 (99% base call accuracy). The high-quality reads (25.5 to 44.0 Million per library, Supplementary Table S2) were mapped to the *Z*. *marina* genome v2.1 (Olsen et al., 2016) with BWA v0.7.17 (Li and Durbin, 2009). Read duplicates were removed with MarkDuplicatesSpark within GATK v4.1.4.1 (Auwera et al., 2013). SNPs were called with HaplotypeCaller, followed by CombineGVCFs and GenotypeGVCFs within GATK v4.1.4.1 (Auwera et al., 2013).

Before filtering SNPs, we excluded indels, non-variant sites, and alternate alleles not present in any genotypes from the vcf file with SelectVariants within GATK v4.1.4.1 (Auwera et al., 2013). This set of 759,407 raw SNPs was reduced to 105,443 SNPs after hard-filtering with vcffilter from vcflib (Garrison, 2020) with thresholds that were based on density plots drawn with ggplot2 (Wickham, 2016): QualByDepth (QD < 15.0), FisherStrand (FS >12.0), RMSMappingQuality (MQ < 38), MappingQualityRankSumTest (MQRankSum < −1.5), ReadPositionRankSumTest (ReadPosRankSumand < −4.0), and Depth (DP > 4000.0, in order to remove SNPs potentially caused by genome duplication). Subsequently, we used VCFtools v0.1.15 to remove genotpes with genotype quality < 30 (--minGQ 30) or depth < 20 (--minDP 20), and to remove SNPs with more than 2 alleles (--min-alleles 2 --max-alleles 2), with a minor allele frequency of 0.01 (--maf 0.01), and with any missing genotype (--max-missing-count 0). From the remaining 15,508 high-quality SNPs (Supplementary File S1) we excluded all that shared the same genotypes among all 10 heat-stressed shoots, as these reflected genetic differences only to the reference genome. The remaining 1,079 SNPs (Supplementary File S2) were used to estimate euclidean genetic distances among the 10 shoots using the R package vcfR v1.9.0 (Knaus and Grünwald, 2017) and the *dist* function of the R package ‘stats’ v3.6.9 (R Core Team, 2019). We tested for correlation between genetic and physical distance among the 10 heat-stressed shoots with Mantel tests in the R package ‘vegan’ v1.4-2 (Oksanen et al. 2019), using 1,000 permutations, and the Pearson’s product moment correlation method.

### 2.7 Methylome characterization

Sequencing libraries were prepared according to the MethylRAD protocol (Wang et al., 2015) with few adjustments. MethylRAD is a genome-reduction method based on the methylation-dependent restriction enzyme FspEI that targets fully methylated CCGG and CCWGG motifs, thus capturing methylation in CG and CHG sequence contexts. MethylRAD has the potential to reveal genome-wide DNA methylation patterns that are consistent with those generated from Whole Genome Bisulfite Sequencing (Wang et al., 2015). First, sense and anti-sense oligos of adapters A1 and A2 (Supplementary File S3) were annealed in 10 μL containing 10 μM of each oligo (Eurofins), 10 mM Tris HCl (Thermo Fisher), 50 mM NaCl (Thermo Fisher), and 1 mM EDTA (Thermo Fisher). Library preparation began with digestion of 100 ng cleaned genomic DNA at 37°C for 4 h in 15 μL containing 4 U FspEI (NEB), 1X CutSmart Buffer (NEB), and 30X Enzyme Activator Solution (NEB). Digestion was verified on a TapeStation 2200 with a D1000 ScreenTape. Secondly, adapters were ligated to the digested fragments over night at 4°C in 26μL containing 13 μL digestion solution, 0.1 μM each of two annealed adapters, 1X T4 ligase buffer (NEB), 1.5 μM ATP (NEB) and 1040 U of T4 DNA ligase (NEB). Ligation products were amplified in 20 μL reactions containing 7 μL ligated DNA, 0.05 μM of each primer (P1 and P2, Supplementary File S3), 0.3 mM dNTP, 1X Phusion HF buffer (NEB) and 0.4 U Phusion high-fidelity DNA polymerase (NEB). PCR was conducted using a Veriti 96-Well Thermal Cycler (Applied Biosystems, Life Technologies) with 16 cycles of 98°C for 5s, 60°C for 20 s, 72°C for 10 s, and a final extension of 5 min at 72°C. The target band (approx. 100 bp) was extracted from a 2% E-Gel (Thermo Fisher). For multiplex sequencing, shoot barcodes were introduced by means of PCR. Each 20 μL PCR reaction contained 12 μL of gel-extracted PCR product, 0.2 μM of each primer (P3 and index primer, Supplementary File S3), 0.3 mM dNTP, 1X Phusion HF buffer (NEB) and 0.4 U Phusion high-fidelity DNA polymerase (NEB). PCR was conducted with the same PCR cycling program outlined above. PCR products were purified using AMPURE XP beads (Beckman Coulter) using a 1.8:1 volume ratio of beads to product, and a final elution in 22 μL EB buffer (Qiagen). The purified fragments were sequenced on an Illumina NextSeq 500 (1×75bp) using a high-output flow-cell.

The sequenced reads were quality-trimmed with TrimGalore! v0.4.1^4^ by removing the adapter sequences with a stringency of 3 bp overlap, low-quality bases with a Phred score Q < 20, and the terminal 2 bp from both ends in order to eliminate artifacts that might have arisen at the ligation position. Quality was checked with FastQC v0.11.8^5^ to control for aberrant read base content, length distribution, duplication and over-representation.

The high-quality reads were mapped with SOAP v1.11 (Li et al., 2008) to 628,255 *in silico* predicted MethylRAD tags that were extracted from the *Z*. *marina* genome v2.1 from ORCAE (Sterck et al., 2012) with the custom python script *InSilicoTypeIIbDigestion.py^6^*. For mapping, we allowed for two mismatches, filtered reads with >1 N, and used a seed size of 8 bp. Based on the uniquely mapped reads, we counted the coverage of each methylated site for each shoot using htseq-count (v0.7.2). Methylation calls were retained only for sites with ≥ 2x coverage, which reduced the false-positive rate from 1.10% to 0.23% for CG sites and from 2.50% to 0.89% for CHG sites. False-positive rates were estimated as the percentage of methylation sites supported by at least two reads in the generally unmethylated chloroplast genome. For each shoot, raw counts were normalized to reads-per-million by dividing reads per site through the total number of reads per shoot library, times one million.

The methylated sites were annotated with the v2.1 gff3 file from ORCAE (Sterck et al., 2012), and separated into genes, intergenic regions, and TEs. TEs were located in both genes and intergenic regions, and contained repeats of the rnd-1/2/3/4 families and some selected repeats (rnd-5-family-7328, rnd-5-family-287, rnd-5-family-4856). The methylated sites were further separated into such containing CG and CHG recognition sites.

### 2.8 Methylation variation between the transect shoots

In order to describe the level of intra- and inter-clonal methylation differences among the 42 transect shoots, we converted the normalized counts to bivariate states (methylated/non-methylated). We then calculated the number of sites with different methylation states pairwise between all 41 ramets of the same genet, and between these ramets and the single other seven-microsatellite-genotype (transect shoot 27, Supplementary table S1).

We tested for correlations between epigenetic and physical distance among the 41 ramets with Mantel tests using the R package ‘vegan’ v1.4-2 (Oksanen et al., 2019) with 1,000 permutations, and the Pearson’s product moment correlation method. *P*-values were adjusted according the Benjamini-Hochberg method (Benjamini and Hochberg, 1995). Epigenetic distance was calculated as the Euclidean distance between shoots based on their coordinates in the 2-dimensional PCA (Principal Component Analysis) plot using the *dist* function of the R package ‘stats’ v3.6.0 (R Core Team, 2019). PCA was done on reads-per-million for all shoots with the PCA function of the R package ‘FactoMineR’ (Lê et al., 2008).

### 2.9 Correlation between methylome variation and photosynthetic performance

For the 10 heat-stressed samples, we tested for correlations between epigenetic distance and performance difference (PiABS values) among shoots while controlling for genetic distance with Partial Mantel tests using the function *partial.mantel.test* of the R package ‘ncf’ v1.2.9 (Bjornstad, 2020). Only significant correlations (p<0.05, adjusted for multiple comparisons according to Benjamini and Hochberg (1995) in R (R Core Team, 2019)) with coefficients *R* > 0.65, when controlled for genetic distance, were considered strong enough to be reckoned as biologically linked. The same analysis ran Mantel tests for correlation between genetic and epigenetic distance, and between genetic distance and performance difference.

### 2.10 Methylome heat stress response

The methylome of the 10 heat-stressed shoots was characterized under control, stress, and recovery conditions (3 shoots died from the stress), and compared with the methylome of field transect samples with PCA on reads-per-million data using the PCA function of the R package ‘FactoMineR’ (Lê et al., 2008). Epigenetic distance between the samples was estimated for each sequence context (gene, intergene, TE, each in CG and CHG regions, respectively) as the Euclidean distance in the 2-dimensional PCA plot using the *dist* function of the R package ‘stats’ v3.6.9 (R Core Team, 2019).

### 2.11 Differential methylation analyses

To estimate the number of sites that changed in methylation state from control to stress and recovery conditions, we applied differential methylation analyses using the R package ‘edgeR’ 3.20.9 (Robinson et al., 2009) within the ‘SARTools’ pipeline v1.6.6 (Varet et al., 2016). Read counts were normalized using a trimmed mean of M-values (TMM) between each pair of samples (Robinson and Oshlack, 2010). *P*-values, indicating the significance of increased or decreased methylation, were adjusted according the Benjamini-Hochberg method (Benjamini and Hochberg, 1995).

To identify how the methylome differed between samples of high and low performance, we used differential methylation analyses using the R package ‘edgeR’ 3.20.9 (Robinson et al., 2009) within the ‘SARTools’ pipeline v1.6.6 (Varet et al., 2016). Read counts were normalized using a trimmed mean of M-values (TMM) between each pair of samples (Robinson and Oshlack, 2010). Differential methylation analysis was done only for conditions and sequence contexts where a positive correlation was found between performance differences and epigenetic distance. For control conditions, we compared methylation levels in genes (CG regions) between the two samples of highest performance (79.1, and 13.2) and the two samples of lowest performance (63.1 and 59.2, Supplementary Figure S1). For recovery conditions, we compared methylation levels in intergenic TEs (CHG regions) between the two samples of highest performance (79.1 and 17.1) and the two samples of lowest performance (57.1 and 59.2).

For differentially methylated sites within gene bodies, we tested with Fisher’s exact tests for enrichment of gene ontology terms of biological processes with the R package ‘topgo’ (Alexa and Rahnenführer, 2010) using Fisher’s exacts tests. GO terms were obtained from the v2.1 *Zostera* genome annotation from the ORCAE database (Sterck et al., 2012). To reduce redundancy in the significantly enriched GO terms (*p*-values < 0.05), we calculated ‘sim rel’ scores (Schlicker et al., 2006) (Allowed similarity=0.5), based on the *A*. *thaliana* GO-term database, using the REVIGO web server (Supek et al., 2011).

## 3 Results

### 3.1 Methylome characterization and variation among the clonal transect shoots

On average, 74 million high-quality reads were obtained per sequencing library, ranging from 0.7 to 151 million (Supplementary Table S3). DNA raw reads are accessible from NCBI under BioProject number PRJNA575339^7^ On average, 35% of the high-quality reads mapped to the *in silico* digested *Z*. *marina* genome, and 11% mapped uniquely (annotated reads-per-million in Supplementary File S4). In total, 144,420 sites were methylated (covered by at least two reads) across all transect shoots. Across all transect shoots, 84,640 methylated CG sites and 59,780 methylated CHG sites were detected, which represents 59% and 41% of all methylated sites, respectively (Supplementary Table S4). About 23% of all methylated sites fell in gene bodies (of which 41% in TEs), and 77% in intergenic regions (of which 42% in TEs) (Supplementary Figure S2). In gene bodies, 67% of the methylated sites fell in CG regions, in intergenic regions only 56%.

Based on their multi-loicrosatellite genotype, 41 of 42 shoots sampled along the transect (except sample 27 in Supplementary table S1) belonged to the same genet (Supplementary Table S5). Methylome variation between these 41 clonal ramets exceeded the methylome shift induced by heat stress (Figure 2), and was not higher between shoots of different genets (Figure 3A,B, Supplementary Table S4). Methylation differences were generally lower in CHG than in CG sequence contexts (Figure 3A,B). Epigenetic distance was not significantly correlated with physical distance in any sequence context (Supplementary Table S6).

**Figure 2.**
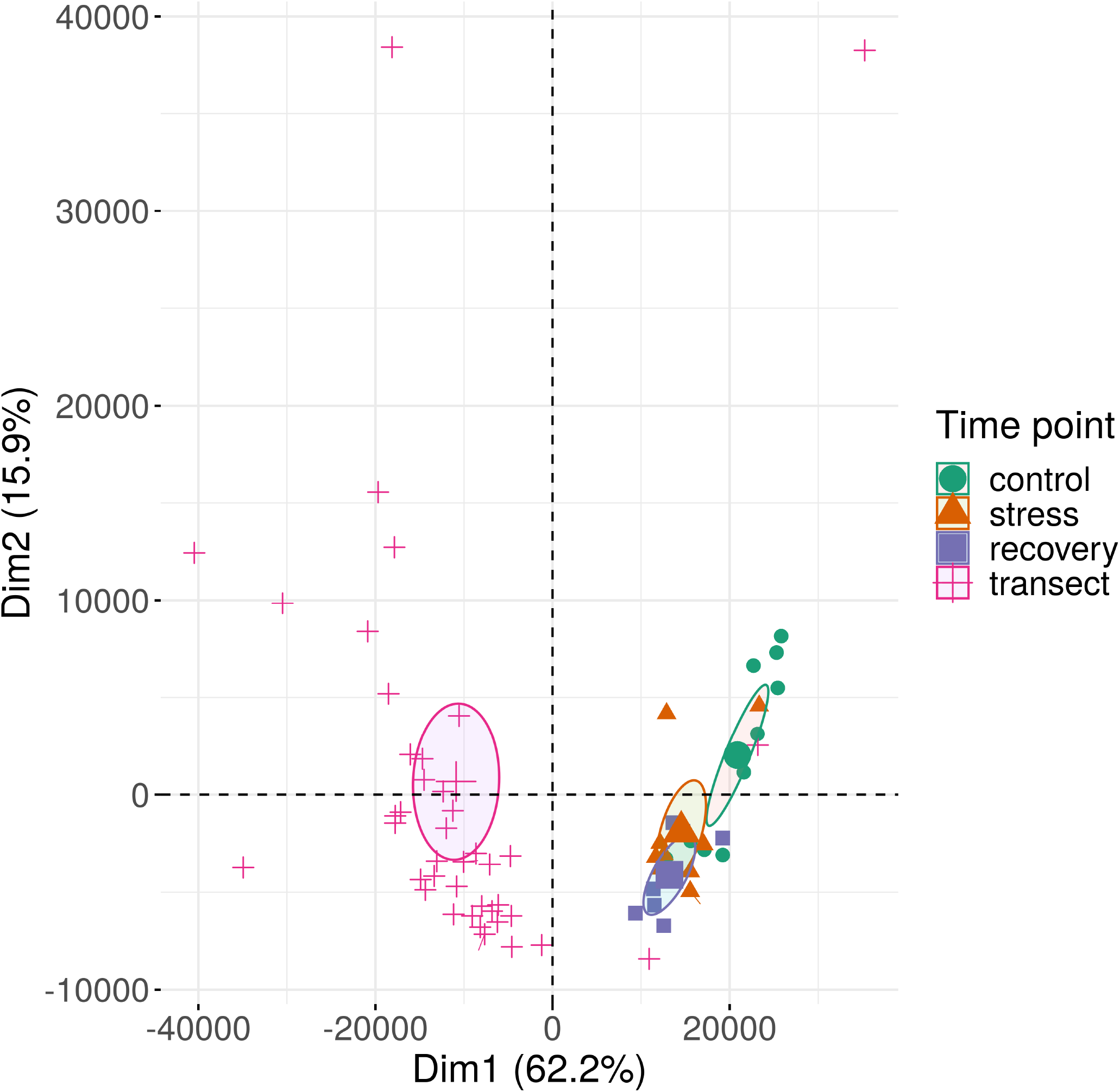
Methylation variation of field transect samples of *Zostera marina* exceeded the shift in methylation patterns along the first two principal components in response to heat stress. The samples are plotted along the first two principle components (Dim) based on methylation profiles across all sequence contexts. Circles represent 95% confidence intervals around group means. Bracketed numbers represent the percentage of explained variation.

**Figure 3.**
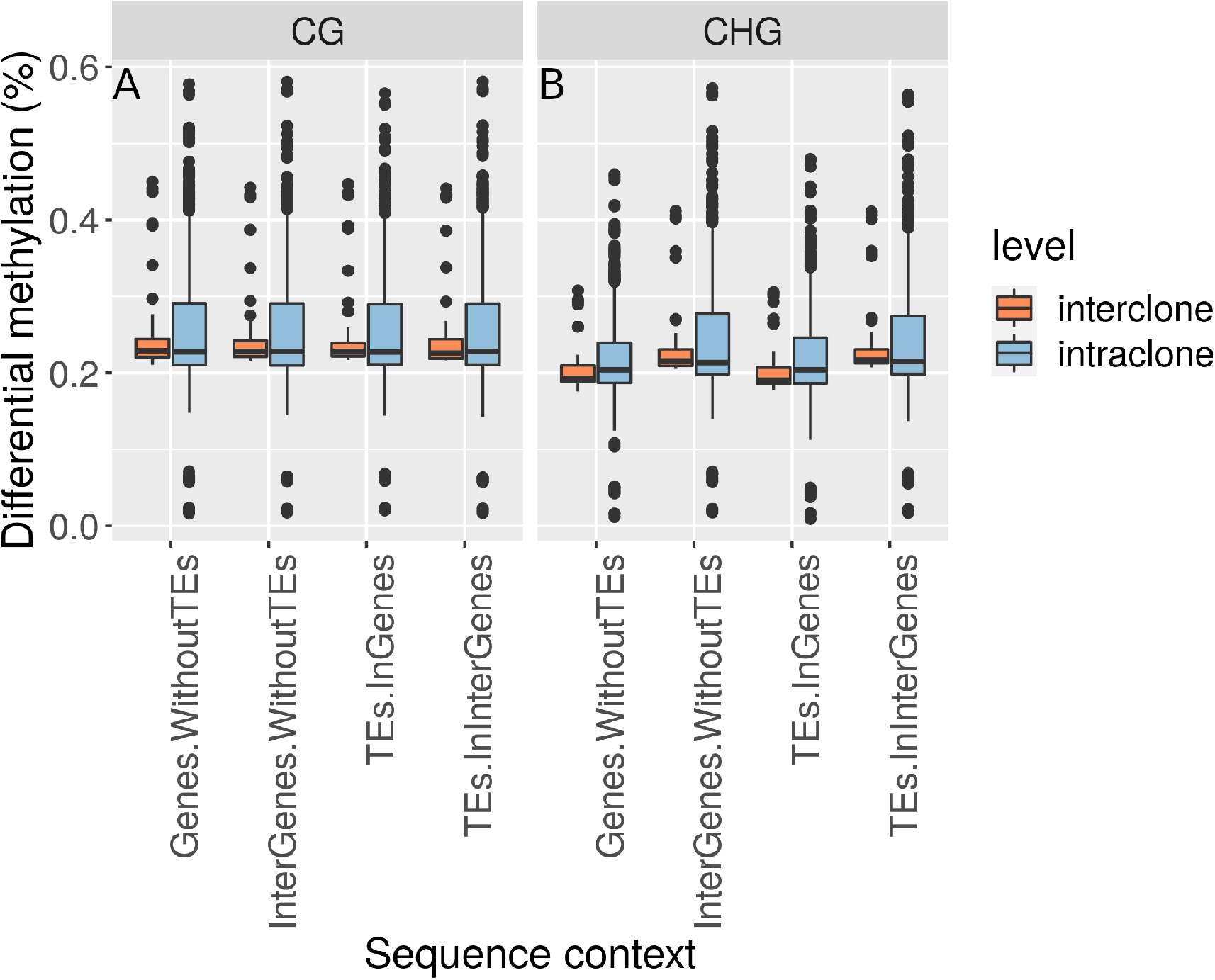
Genetic relatedness did not restrict epigenetic differentiation. Bivariate methylation differences in (A) CG and (B) CHG regions were comparable within ramets of the same genet (intraclone) and between ramets of different genets (interclone) of *Zostera marina* across all sequence contexts.

### 3.2 Genetic variation among heat-stressed shoots

Based on whole genome sequences of the 10 heat-stressed shoots, we identified 15,508 high-quality SNPs (Supplementary File S1). That all 10 shoots shared the same heterozygous state in 14,431 (93%) of all SNPs, and shared the same multi-locus microsatellite genotype (Supplementary Table S5), suggests that they were clones having originated from one ancestral zygote (Supplementary Figure S3) (Yu et al., 2020).

Despite clonal, the 10 shoots differed in 1.079 SNPs resulting from somatic mutations. Based on these SNPs, euclidean genetic distances ranged from 9 to 35 (frequency distribution in Supplementary Figure S4), and were not significantly correlated with physical distances between shoots (*R*=0.18, *p*=0.12).

### 3.3 Link between epigenetic distance and performance changes under heat stress

Photosynthetic performance (PiABS) declined in all 10 shoots under heat stress (Figure 1, Supplementary Table S7). Seven of the 10 shoots recovered from the heat stress, yet performances did not reach pre-stress levels. Performances under control and recovery conditions were positively correlated (adjusted *p*<0.05, *R*=0.93, Figure 4).

**Figure 4.**
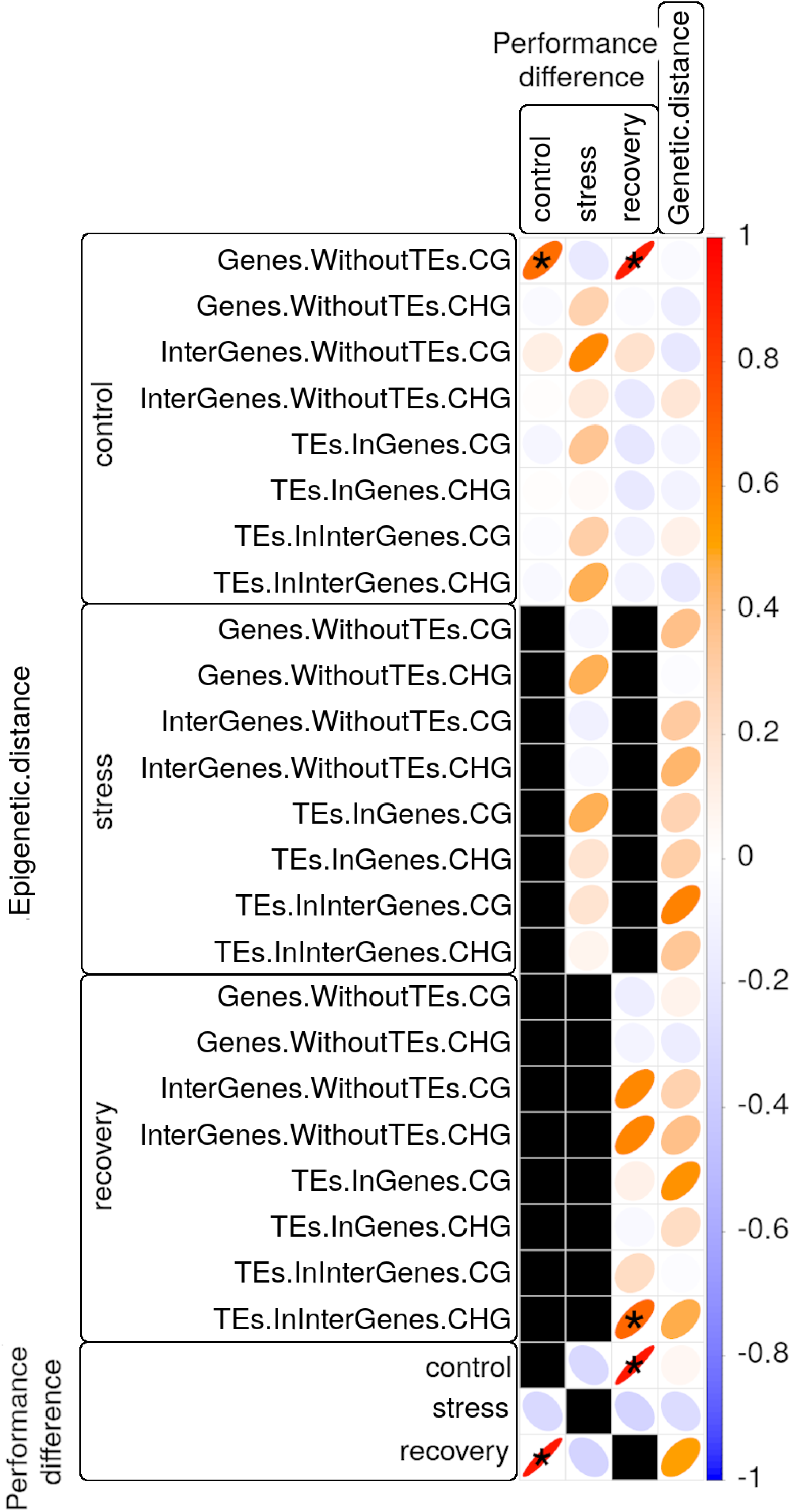
Correlation matrix of photosynthetic performance (PiABS) difference (first three columns) or of genetic distance (last column) among clonal *Zostera marina* shoots with epigenetic distance (first 24 rows) or with performance difference at control, stress, or recovery conditions (last three rows). All correlations between performance difference and epigenetic distance were controlled for genetic distance. Black squares represent untested correlations that were considered biologically not meaningful. Pearson product-moment correlation coefficients R are encoded by the color gradient explained in the bar on the right end, and by the shape of the ellipses. Narrow ellipses represent stronger correlations as compared with wide ones. Asterisks highlight strong (*R*>0.65) and significant (adjusted *p*<0.05) correlations. CG and CHG: sequence contexts of the methylated cytosine; TE: Transposable element.

Genetic distance correlated moderately (4<*R*<6, adjusted *p*<0.05) with performance differences in recovered samples, and with epigenetic distance in some sequence contexts of stressed and recovered samples (Figure 4). Nevertheless, epigenetic distance correlated strongly (*R*>0.65, adjusted *p*<0.05) with performance differences even after controlling for genetic distance based on 1,079 SNPs: 1) epigenetic distance among control samples in CG gene body regions correlated with performance differences prior to stress, and after recovery (stress resilience, Figure 4); epigenetic distance among recovered samples in CHG regions of intergenic TEs correlated with performance differences after recovery (Figure 4).

### 3.4 Methylome heat stress response

Methylation patterns in all sequence contexts changed under heat stress and did not return to but instead diverged further from control (pre-stress) patterns during the recovery period (Figure 5 for all sequence contexts combined, annotated reads-per-million in Supplementary File S5, Supplementary figure S5 showing the methylation shift for the different sequence contexts). More sites became hyper-than hypo-methylated in response to stress (Figure 6A,D). Methylation levels differed significantly at 437 sites between control and stress conditions (257 hyper- and 180 hypo-methylated), at 1788 sites between control and recovery conditions (1141 hyper-, and 647 hypo-methylated), and only at 39 sites between stress and recovery conditions (18 hyper-, and 21 hypo-methylated, Supplementary Table S8). After recovery, CG methylation had been more heat-responsive in gene body regions, and CHG methylation in intergenic regions (Figure 6B,E).

**Figure 5.**
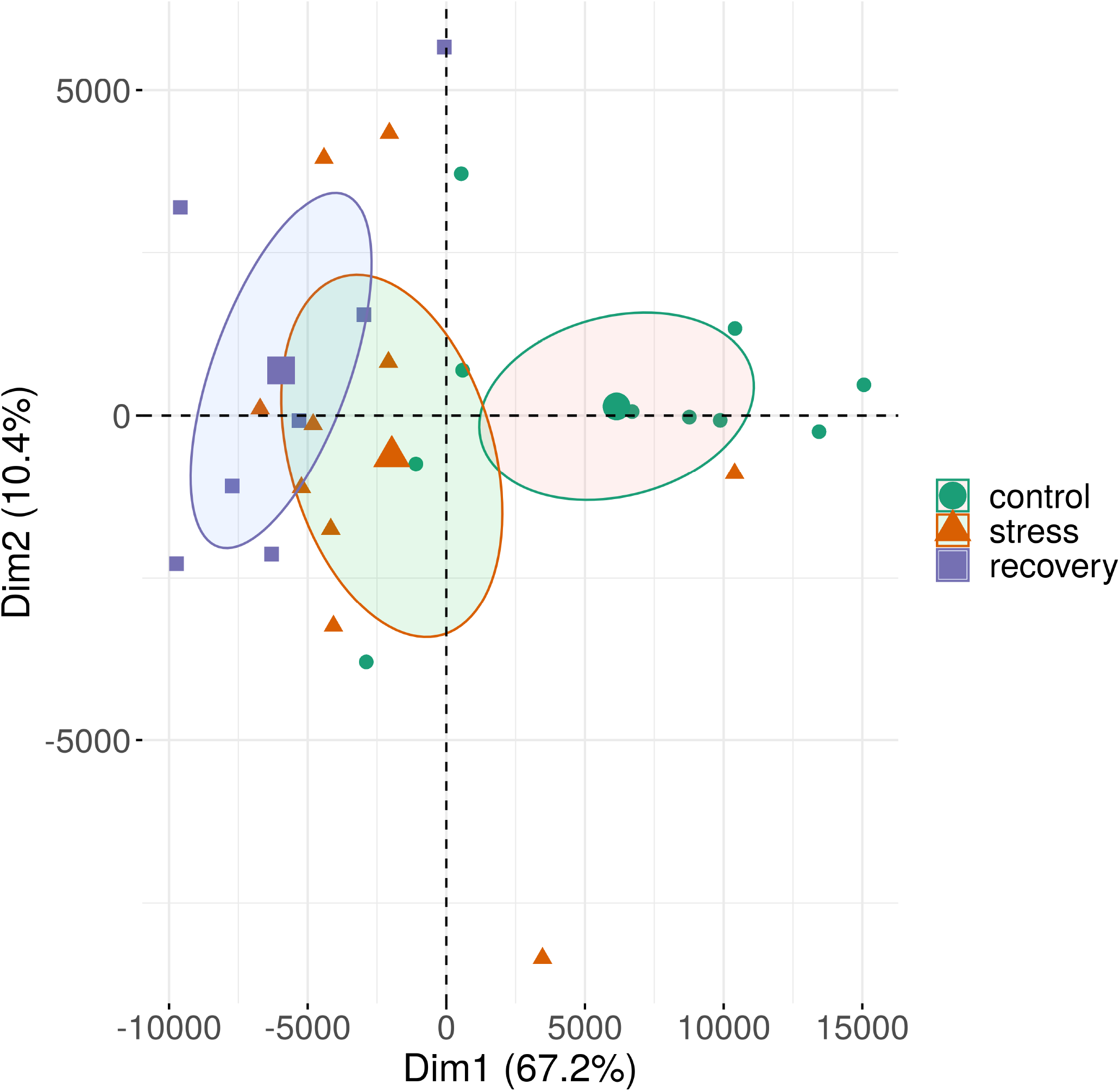
Methylation patterns of all sequence contexts in shoots of a *Zostera marina* clone changed in response to heat stress and did not return to pre-stress patterns after a >5-week recovery period. See Supplementary Figure S5 for methylation shifts separated by sequence context. The samples are plotted along the first two principle components (Dim) based on methylation profiles across all sequence contexts. Circles represent 95% confidence intervals around group means. Bracketed numbers represent the percentage of explained variation.

**Figure 6.**
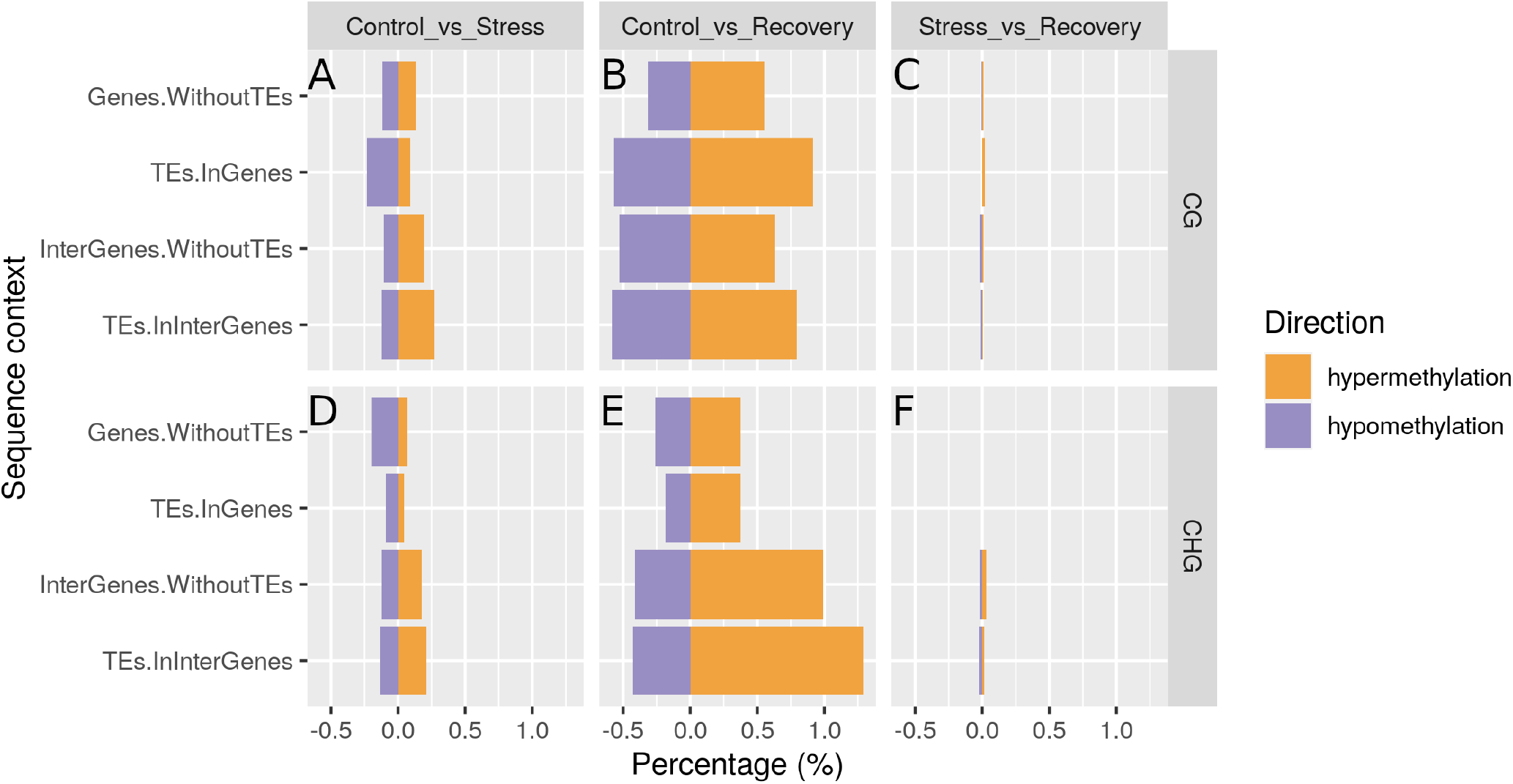
Hyper-methylation dominates the stress response of *Zostera marina*. Hyper- and hypomethylated sites in CG (A-C) and CHG (D-F) regions in stressed vs. control samples (A,D), recovered vs. control samples (B,E), and in recovered vs. stressed samples (C,F). Stress-responsive methylation changes were memorized over the >5-week recovery period. TE: transposable element.

After recovery, methylation increased in gene bodies with functions including DNA transcription and replication; catabolism of misfolded proteins, gamma-aminobutyric acid (GABA), and neurotransmitter; as well as amino acid synthesis (Supplementary Table S9). DNA methylation had decreased in gene bodies with functions including transmembrane transport of cations, ions, and protons; transport of ammonium and phospholipids; localization of lipids and organelles; exocytosis; and secretion (Supplementary Table S9).

### 3.5 Differential methylation between shoots of high and low photosynthetic performance

Recovered shoots of high and low performance presented highly similar methylation patterns. Only seven CHG sites in intergenic TEs were hyper-methylated in recovery samples of high photosynthetic performance (Supplementary Table S10). In contrast, 89 CG gene body sites were hyper-methylated (1 hypo-methylated) in control samples of high photosynthetic performance (Supplementary Table S10). Enriched biological processes in the 89 hyper-methylated CG sites included ‘light harvesting in photosystem I’ and ‘protein folding’ (Supplementary Table S11).

## 4 Discussion

Plant genets persisting >1,000 years challenge the positive correlation between genetic variation retained through recombination, and long-term survival. Our study shows for the first time that ramets of the same seagrass genet display DNA methylation variation that is associated with phenotypic variation in the fitness-related traits of photosynthetic performance and heat stress resilience. Heat responsive methylation changes were memorized for at least five weeks in the recovered shoots. This supports the hypothesis (Dodd and Douhovnikoff, 2016) that methylation variation, via variation in gene regulation, compensates potential costs of clonal reproduction (Lynch et al., 1993; Lynch and Lande, 1998), and contributes to the long-term survival of clonal seagrass meadows by increasing temporally stable variation in ecologically relevant traits that can not be simply explained by the underlying somatic genetic variation.

### 4. Methylome variation of functional relevance

The seagrass methylome is flexible and responds directly to environmental change, given its strong change from field transect samples to acclimated control samples within two weeks (Figure 2). Methylome variation among ramets of the same genet resulted either from random epimutations or from microscale variations in the environment because its correlation with geographic distance between shoots was weak. The change in depth of 3 meters and, thus, gradual changes in environmental factors along the sampled transect may not have been extreme enough to impose a result. The disagreement between methylome variation and transect position may also result from recent uprooting and re-settling of some shoots. This was suggested to explain disagreement between genetic similarity of clonal shoots and their transect position in another clonal meadow of *Z*. *marina* (Yu et al., 2020), and would further explain the absence of correlation between genetic and physical distance in our heat-stressed ramets. In such case, the re-settled shoots would display a methylome shaped by distant environmental conditions. Whether environmentally induced or not, methylome variation in CG gene regions predicted pre- and post-stress performance (Figure 4) and, thus, is likely functionally relevant for thermal acclimation and stress resilience.

Although difficult to prove (Crisp et al., 2016), the correlations between methylome and photosynthetic performance differences appear to be causal, given that samples of high performance showed hyper-methylated CG sites in gene bodies with relevant functions: ‘light harvesting in PSI’, ‘protein folding’, ‘protein refolding’, and ‘misfolded or incompletely synthesized protein catabolic process’ (Supplementary Table S11). These functions can prime against heat stress when increased methylation is associated with expression of the underlying genes and, thus, with accumulation of protective molecules like heat-shock proteins that are involved in repair and prevention of heat-induced protein damage (Feder and Hofmann, 2002), and that can facilitate a fast stress response. Indeed, while genes are generally repressed by methylated promoter regions (Lisch, 2009), genes are often activated by CG gene body methylation (Zhang et al., 2006; Schmitz et al., 2013; Yang et al., 2014; Dubin et al., 2015; Bewick and Schmitz, 2017; Niederhuth and Schmitz, 2017) that prevents aberrant expression from intragenic promoters (Takuno and Gaut, 2012; Neri et al., 2017). Thus, via association with gene expression, the observed methylome variation can putatively create ecologically relevant phenotype variation.

Although our study was based on clonal ramets that presented identical seven-locus microsatellite genotypes, the 10 heat-stressed ramets varied genetically at 1,079 SNPs generated by somatic mutations. Epigenetic differentiation between populations can be tightly linked to genetic differentiation (Herrera and Bazaga, 2010; Gáspár et al., 2019). Accordingly, trans-acting SNPs appeared to explain the correlation between gene body methylation and latitude in *Arabidopsis* (Dubin et al., 2015). In contrast, part of the correlation between DNA methylation with ecological factors in landscape epigenomic studies on non-model plant species could not be predicted from the observed underlying patterns of genetic relatedness (Schulz et al., 2014; Foust et al., 2016; Gugger et al., 2016). Genetic variation could also not explain the association between epigenetic variation and phenotypic variation (leaf, petiole and functional traits) in natural populations of the perennial herb *Helleborus foetidus* (Medrano et al., 2014), and in clonal populations of the introduced clonal herb *Hydrocotyle vulgaris* (Wang et al., 2020). In agreement, the correlation between methylome variation and stress performance/recovery in our study could not be predicted from the underlying genetic variation. Independent from genetic variation was also the methylome stress-response that provides potential for temporally stable phenotypic change at time scales unattainable by somatic mutations. Thus, our results provide a first indication that DNA methylation variation provides a layer of ecologically relevant, and potentially selectable, phenotypic variation that is independent from genetic variation in clonal seagrass meadows.

### 4.2 Potential of the epigenetic heat stress response for heat-hardening

Stress recovery is a critical period during which the degree of resetting versus memory of epigenetic stress response is determined (Crisp et al., 2016). *Z*. *marina* memorized much of the methylome stress response after a 5-week recovery period (Figure 5). The methylation changes from control to recovery conditions were initiated in the stress phase, and then continued to diverge from control conditions. This is illustrated by the strong similarities in methylation profiles between the stress and recovery phase, with an apparently stronger divergence of recovery profiles from the control ones (Figure 6B,E, Supplementary Table S8). This agrees with increasing divergence of gene transcription profiles from heat-stress to recovery conditions in a Danish *Z*. *marina* population (Franssen et al., 2011). We speculate that gene expression changes, and other molecular mechanisms involved in the heat-stress response, could have triggered additional methylation changes after the stress was removed (e.g. Secco et al. 2015) The memory of the epigenetic heat stress response is long enough to potentially heat-harden the same generation of previously exposed shoots.

In agriculture, hardening/priming of seeds is a long-standing practice to enhance crop resistance to environmental challenges, including hot, cold, dry, or saline conditions, or pathogen infections (Ibrahim, 2016; Wojtyla et al., 2016; Pawar and Laware, 2018). Heat-priming has only very recently been described in seagrass (Nguyen et al., 2020). *Zostera muelleri* and *Posidonia australis* both performed better under a second heat-wave, in terms of photosynthetic capacity, leaf growth, and chlorophyll a content, when they had been previously exposed to a first heatwave as compared with naïve controls (Nguyen et al., 2020). This could explain why no mortality was reported for the seagrass *Posidoinia oceanica* after intense and log-lasting heat-waves in 2012, 2015, and 2017 (Darmaraki et al., 2019), although it had suffered high mortality rates after the 2006 heatwave (Marbà and Duarte, 2010), as discussed in Nguyen et al. (2020).

Molecular mechanisms involved in forming a stress memory include stalled RNA polymerase II, storage of chemical signaling factors, accumulation and phosphorylation of transcription factors, and epigenetic mechanisms such as microRNAs, histone modifications, and DNA methylation (Iwasaki and Paszkowski, 2014; Crisp et al., 2016; Hilker et al., 2016; Gallusci et al., 2017; Lämke and Bäurle, 2017). In heat-primed seagrass, the significant regulation of methylation related genes, in particular histone methyltransferases, suggests that epigenetic modifications contribute to form a thermal stress memory (Nguyen et al., 2020). Our study adds evidence that also DNA methylation is likely involved in this process. While the memory of histone modifications lasts generally not longer than hours or days (Cedar and Bergman, 2009; Huang et al., 2013; Lämke and Bäurle, 2017; Kumar, 2018), DNA methylation modifications have been shown to be partly transgenerationally stable under both sexual (Ou et al., 2012; Herman and Sultan, 2016), and asexual reproduction (Raj et al., 2011; Verhoeven and van Gurp, 2012; González et al., 2017). For example, that DNA methylation memories can mediate adaptive transgenerational plasticity was shown in the plant *Polygonium persicaria*, in which offspring demethylation with zebularine removed the adaptive effect of parental drought in form of longer root systems and greater biomass (Herman and Sultan 2016).

Across mitotically grown generations, epigenetic patterns can be expected to be more faithfully inherited than across sexual generations, because clonal growth circumvents epigenetic reprogramming during meiosis and embryogenesis (Hirsch et al., 2012; Douhovnikoff and Dodd, 2014; Verhoeven and Preite, 2014; Crisp et al., 2016; Latzel et al., 2016). CG methylation, maintained by a homologue of mammalian DNA METHYLTRANSFERASE (MET1) (Law and Jacobsen, 2010), is likely more stable than CHG methylation, which is maintained by a dynamic process involving plant-specific CHROMOMETHYLASE and a feed-forward loop involving histone 3 lysine 9 methylation (Law and Jacobsen, 2010). Thus, we propose that memorized changes in CG gene body methylation (Figure 6B), are the most likely methylation marks that may contribute to stress priming with potential for transgenerational plasticity across mitotically grown seagrass generations. Under global warming, however, seagrasses may increasingly investigate in sexual reproduction (Ruiz et al., 2018; Marín-Guirao et al., 2019) and, thus, may be able to transmit only methylome-based stress memories that escape epigenetic reprogramming.

Heat-responsive DNA methylation changes in plants appear not to show a consistent trend across different species, and little is yet known about their functional role (Liu et al., 2015). In *Brassica rapa*, heat-responsive DNA methylation was shown to be associated with differential expression of genes involved in heat stress signal transduction and in RNA metabolic processing (Liu et al., 2018). In *Z*. *marina,* biological functions affected by stress-responsive methylation changes suggest that the plants were investing after stress in the breakdown and repair of misfolded proteins, synthesis of new proteins, growth (DNA replication), and the regulation of transmembrane transport of ions and protons, and neurotransmitter levels (Supplementary Table S9). Of particular interest is the CG hyper-methylation and, thus, potentially constitutive upregulation (Zhang et al., 2006; Schmitz et al., 2013; Yang et al., 2014; Dubin et al., 2015; Niederhuth and Schmitz, 2017) of genes involved in the catabolism of misfolded proteins (Supplementary Table S9). Increased investment in the breakdown of heat-denatured proteins (Feder and Hofmann, 2002), may prime/adapt the affected shoots to future heat stress. This function was also hyper-methylated in control samples of highest performance and stress resilience (Supplementary Table S11). Experimental removal of DNA methylation, e.g. using zebularine or 5-Azacytidine (Griffin et al., 2016), or the targeted change of methylation patterns, e.g. via CRISPR (Xu et al., 2016), will ultimately allow to identify the relationship between heat-responsive methylation patterns and adaptive phenotypic changes.

### 4.3 Conclusions and Perspectives

Our study suggests that DNA methylation is functionally relevant for photosynthetic performance and heat stress resilience, independent from underlying somatic mutations. A portion of the stress-responsive changes in methylation appears sufficiently stable (Figure 5) to allow for medium to long-term acclimation/heat hardening in clonal lineages. In seagrass meadows composed of several genotypes, stress resilience, growth, and associated invertebrate species diversity is enhanced by genotypic variation (Hughes and Stachowicz, 2004; Reusch et al., 2005; Ehlers et al., 2008). In clonal meadows, epigenetic variation may play a similar role in the potential to secure function and resilience not only of *Z*. *marina* plants, but also of the entire associated ecosystem.

Due to anthropogenic stressors, nearly one-third of global seagrass area has disappeared over the last 100 years, and the rate of loss accelerated from ca. 1% yr^−1^ before 1940 to 7% yr^−1^ since 1990 (Waycott et al., 2009). At the same time, rising temperatures open up new thermally suitable habitat in the Arctic (Krause-Jensen and Duarte, 2014). How fast and far warm-temperate and subarctic range edges will move polewards depends on the ability of seagrass to rapidly acclimate and adapt to rising temperatures and other environmental changes (Duarte et al., 2018). Thus, future studies are needed to assess the adaptive value and transgenerational stability of the epigenetic stress response, and to compare the ability to build up epigenetic variation between seagrass meadows of composed of a single or multiple clones, as well as between range center *versus* edge populations.

The functional role of methylation variation in plant genets is not only of fundamental interest but also of applied interest for management programs of clonal organisms designed to assess evolutionary potential and population stability, and to minimize the loss of biodiversity. Our results can be relevant for restoration of seagrass ecosystems that largely depends on the success of replanted shoots to overcome natural variability/stress (Suykerbuyk et al., 2016; van Katwijk et al., 2016). Together with Nguyen et al. (2020) our study suggests that heat priming may improve restoration success of meadows damaged after heat waves (e.g. Arias-Ortiz et al. 2018). Given that 40% of all plant species can reproduce clonally (Tiffney and Niklas, 1985), our findings are further important to other fields, such as invasion biology and crop breeding strategies (Bilichak and Kovalchuk, 2016) of clonal plants.

## 6 Acknowledgements

We thank Marvin Choquet from Nord University for help with re-planting the shoots in the wet lab. We acknowledge Steinar Johnsen from Nord University for setting up the aquaria and water flow in the wet lab. This work was supported by the Norwegian Research Council (Havkyst project 243916), the Åbo Akademi University Foundation sr to CB, and a personal research talents grant from Nord University to AJ. Open access publication fees were covered by Nord University’s Opean Access fund. The content of this manuscript has previously appeared online as a preprint in bioRxiv (Jueterbock et al., 2019).

## 7 Author Contributions

GH (project leader) and AJ were planning the project and designing the experiments. AJ, CB, IS, and GH collected the shoots. AJ analysed the data and wrote the manuscript. AJ, MK, AKSD, JAC, and IS performed the DNA extraction, library preparation, and sequencing. SAH, JLO, and YVdP were involved in data interpretation. All co-authors read and commented on the manuscript.

## Footnotes

1 Available upon publication

2 http://www.bioinformatics.babraham.ac.uk/projects/fastqc/

3 https://www.bioinformatics.babraham.ac.uk/projects/trim\_galore/

4 https://www.bioinformatics.babraham.ac.uk/projects/trim\_galore/

5 http://www.bioinformatics.babraham.ac.uk/projects/fastqc/

6 http://marinetics.org/2017/04/11/REdigestions.html

7 Available upon publication

