## Supplementary material captions for "The seagrass methylome memorizes heat stress and is associated with variation in stress performance among clonal shoots"

### 1 Supplementary Figures

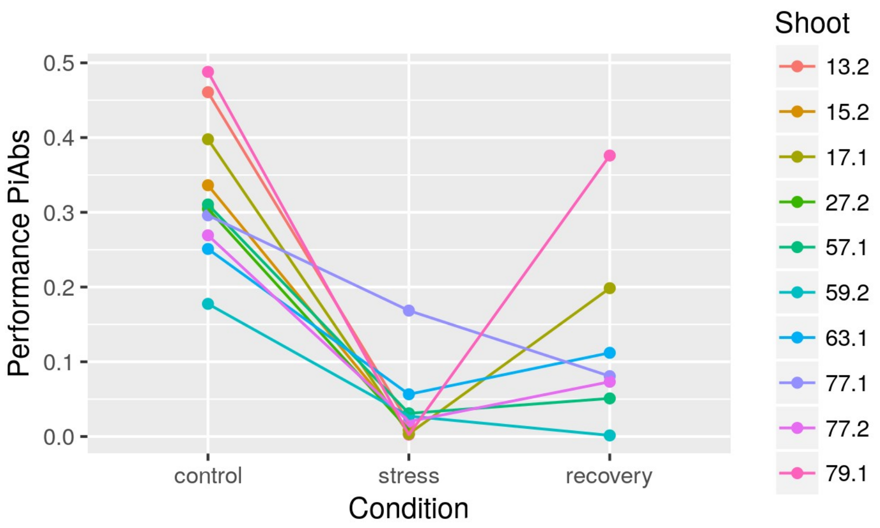


**Supplementary Figure S1** Photosynthetic performance (PiAbs, absolute values) for all ten heat-stressed *Zostera marina* shoots at control, stress, and after recovery. Shoots 13.2, 15.2, and 27.2 did not recover from the stress.

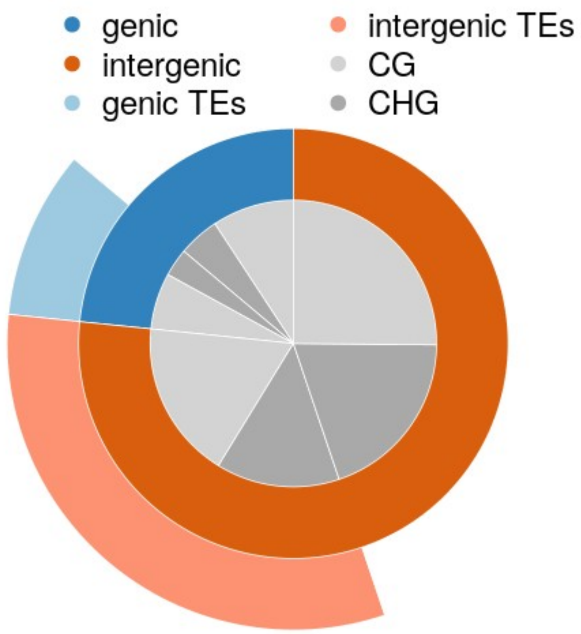


**Supplementary Figure S2** Proportion of methylated sites detected in genes, intergenic regions, and transposable elements (TEs), separated by CG and CHG sequence contexts.

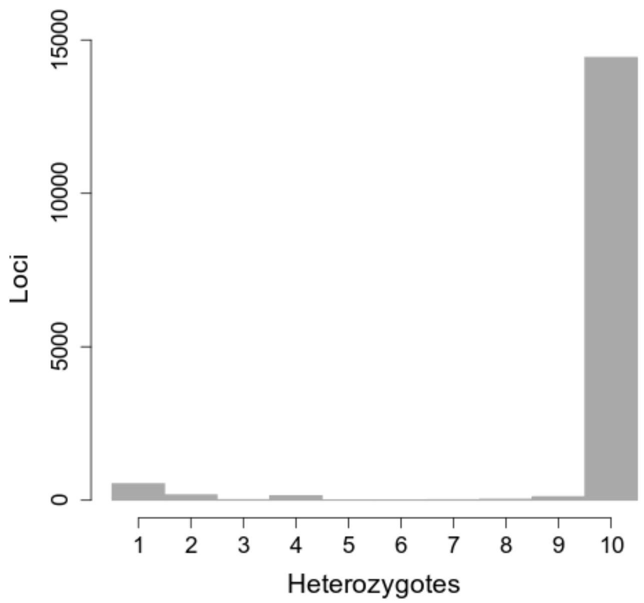


**Supplementary Figure S3** Number of heat-stressed *Zostera marina* shoots sharing the same heterozygous state for 15,508 biallelic SNPs. The dominant peak of loci where all shoots are identically heterozygous suggests that all ten shoots belong to the same genet, and inherited the heterozygous state

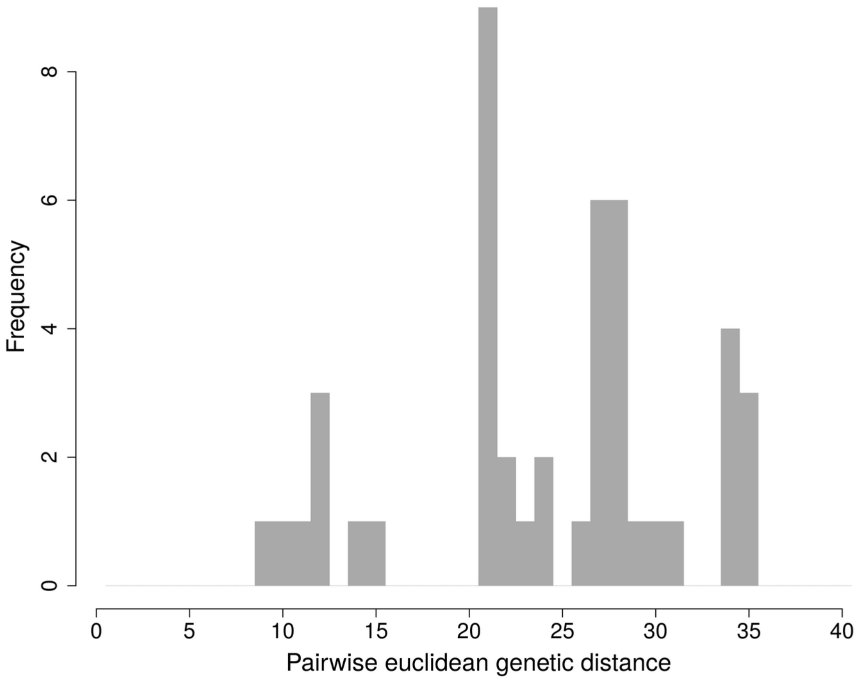


**Supplementary Figure S4** Frequency distribution of pairwise euclidean distances among 10 heat-stressed *Zostera marina* shoots based on 1,079 SNPs resulting from somatic mutations.

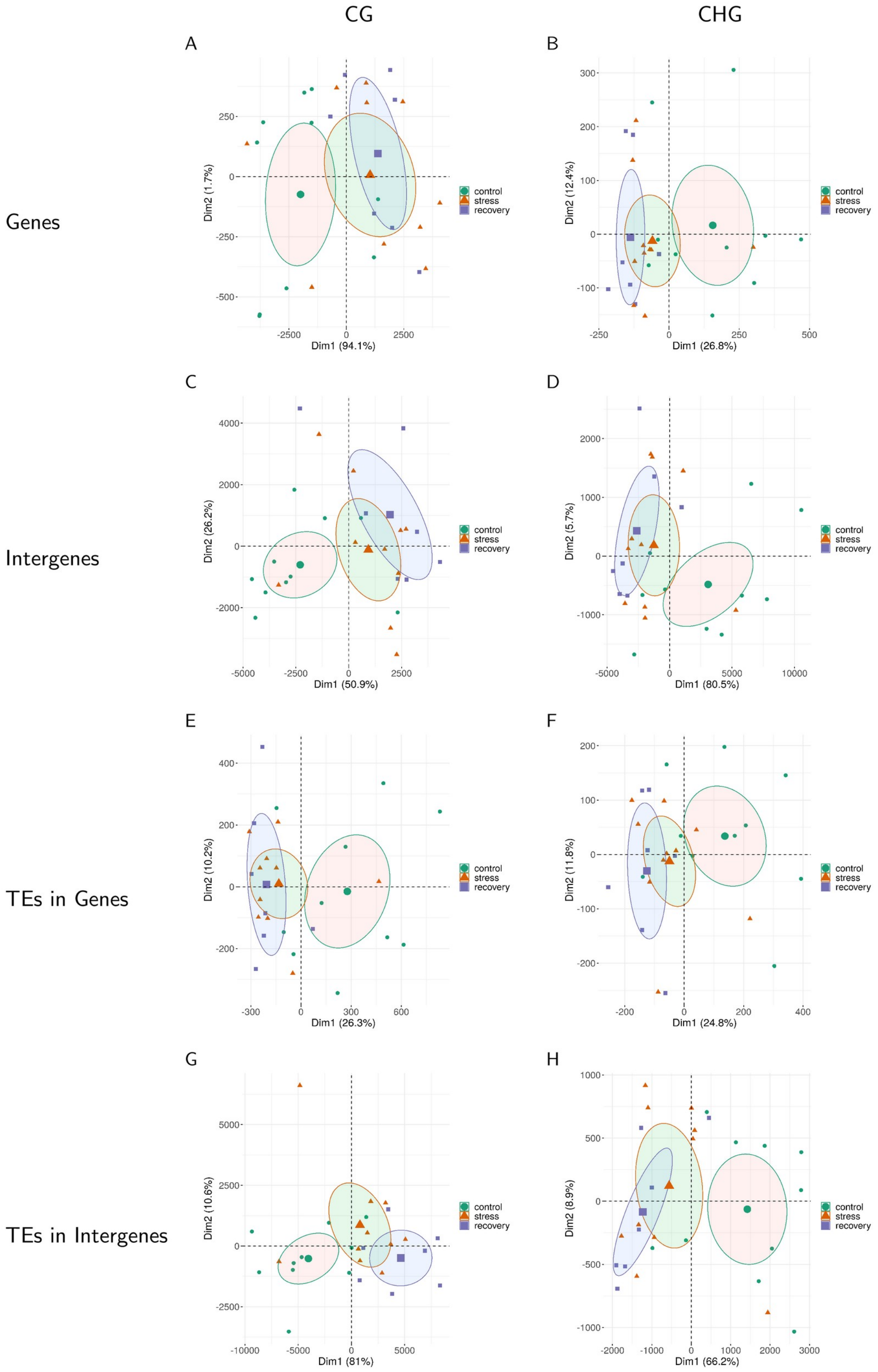


**Supplementary Figure S5** Methylation patterns in shoots of a *Zostera marina* clone changed in response to heat stress and did not return to pre-stress patterns after a 5-week recovery period. This applies for all sequence contexts, including genes, intergenes, and transposable elments (TEs) in genes or intergenes in either CG or CHG sequence contexts The samples are plotted along the first two principle components (Dim) based on methylation proFiles across all sequence contexts. Circles represent 95% confidence intervals around group means. Bracketed numbers represent the percentage of explained variation.

### 2 Supplementary Tables

**Supplementary Table S1** Seagrass samples that were taken along a ca. 250 m transect. Listed are the location and IDs of the 42 samples that were sequenced directly after sampling, and of the 10 samples that took part in the heat stress experiment.

**Supplementary Table S2** Number of raw and filtered (high-quality) reads of genomic libraries prepared for each of the ten heat-stressed samples according to the TruSeq DNA PCR-Free (Illumina) protocol, and sequenced on one Illumina HiSeq 3/4000 lane (2x150bp).

**Supplementary Table S3** Number of sequenced reads, trimmed (high-quality) reads, mapped, and uniquely mapped reads for each sample.

**Supplementary Table S4** Number of methylated CG and CHG sites in genes, intergenic regions, and transposable elements, listed for each transect sample. The minimum, mean, and maximum number of regions that were differently methylated between the samples are listed in the last two columns. Comparisons with sample 27 are interclonal comparisons, the other intraclonal.

**Supplementary Table S5** Microsatellite raw data. Microsatellite fragment lengths for 7 markers (2 columns = 2 alleles per marker) for each sample (ID in the first column).

**Supplementary Table S6** Mantel test results (p-values adjusted by Benjamini-Hochberg correction and Pearson's Correlation Coefficient R) for correlations between epigenetic and geographic distance at each sequence context.

**Supplementary Table S7** Photosynthetic performance measures. Mean values and standard errors of PiABS for each of the 10 shoots at all three sampling points in the heat stress experiment.

**Supplementary Table S8** Number of differentially methylated sites between control vs. stress, control vs. recovery, and stress vs. recovery samples listed for each sequence context.

**Supplementary Table S9** Enriched biological processes showing differential methylation in gene bodies between control and recovery samples. The Supplementary Table lists the direction of methylation (hyper- or hypomethylated in comparison with control samples; whether the methylation site falls within a transposable element; the sequence context (CG or CHG); the GO term ID; description; frequency (proportion of this GO term in the underlying *Arabidopsis thaliana* protein annotation database; log10 p-value of the enrichment test; the term's uniqueness; dispensability (the semantic similarity threshold at which the term was removed from the list and assigned to a cluster); the representative GO term; and a boolean indicating whether the ID stands alone or was assigned to a cluster (eliminated).

**Supplementary Table S10** Differentially methylated sites between two samples of highest (79.1 and 13.2 for control conditions, 79.1 and 17.1 for recovery conditions) and two samples of lowest photosynthetic performance (63.1 and 59.2 for control conditions, 57.1 and 59.2 for recovery conditions). The Supplementary Table contains the following columns: Id, unique MethylRAD tag identifier (location in the genome v2.1); sampleName, raw counts per sample; norm.sampleName, rounded normalized counts per sample; baseMean, base mean over all samples; lowperformance and highperformance, means (rounded) of normalized counts of the biological conditions; FoldChange, fold change of expression; log2FoldChange, estimated by the GLM model. It reflects the differential expression between Test and Ref. If this value is around 0 (the feature methylation is similar in both conditions), positive (the feature is more methylated in the samples of high photosynthetic performance), negative, (the feature is less methylated in the samples of high photosynthetic performance); pvalue, raw p-value from the statistical test; padj, p-value adjusted by Benjamini-Hochberg correction; tagwise.dispersion, dispersion parameter estimated from feature counts; trended.dispersion, dispersion parameter estimated with splines; GeneID, gene identifier based on the /Zostera marina/ genome annotation v2.1; Gene Name; GO Terms.

**Supplementary Table S11** Enriched biological processes with increased methylation in control samples of high performance. The Supplementary Table lists the GO term ID; description; frequency (proportion of this GO term in the underlying *Arabidopsis thaliana* protein annotation database; log10 p-value of the enrichment test; the term's uniqueness; dispensability (the semantic similarity threshold at which the term was removed from the list and assigned to a cluster); the representative GO term; and a boolean indicating whether the ID stands alone or was assigned to a cluster (eliminated).

### 3 Supplementary Files

**Supplementary File S1** Biallelic Single Nucleotide Polymorphisms (15,508) that differ from the reference genome or among the 10 heat-stressed samples of the seagrass *Zostera marina*.

**Supplementary File S2** Biallelic Single Nucleotide Polymorphisms (1,079) that differ among the 10 heat-stressed samples of the seagrass *Zostera marina.*

**Supplementary File S3** *Adapters and primers used to prepare MethylRAD libraries.*

**Supplementary File S4** Annotated reads-per-million for 42 transect shoots based on the Z. *marina* genome annotation v2.1 (GenBank Accession nos. LFYR00000000) listing the geneID, location on the Scaffold, reads-per-million for all samples, region (genic or intergenic), TE (whether it falls within a transposable elment), Name (gene name), GOs (associated gene ontology terms), and SequenceContexts (CG or CHG). The data are openly available in figshare at https://figshare.com/s/6ae9a7224af71be9527b (will be replaced by DOI upon publication).

**Supplementary File S5** Annotated reads-per-million for experimental samples (control, stress, and recovery) based on the *Z. marina* genome annotation v2.1 (GenBank Accession nos. LFYR00000000) listing the geneID, location on the Scaffold, reads-per-million for all samples, region (genic or intergenic), TE (whether it falls within a transposable elment), Name (gene name), GOs (associated gene ontology terms), and SequenceContexts (CG or CHG). The data are openly available in figshare at https://figshare.com/s/75da19b406d1a8d776a7 (will be replaced by DOI upon publication).
