## Supplementary figures and images for "The seagrass methylome memorizes heat stress and is associated with variation in stress performance among clonal shoots"

### Supplementary Figure S1

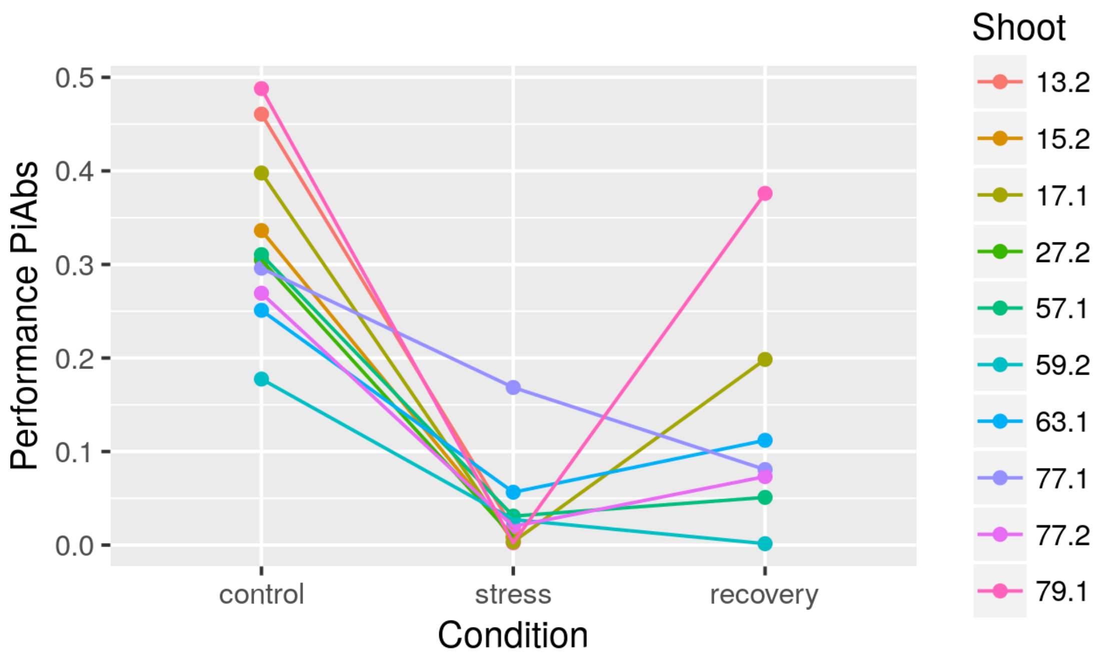

### Supplementary Figure S2

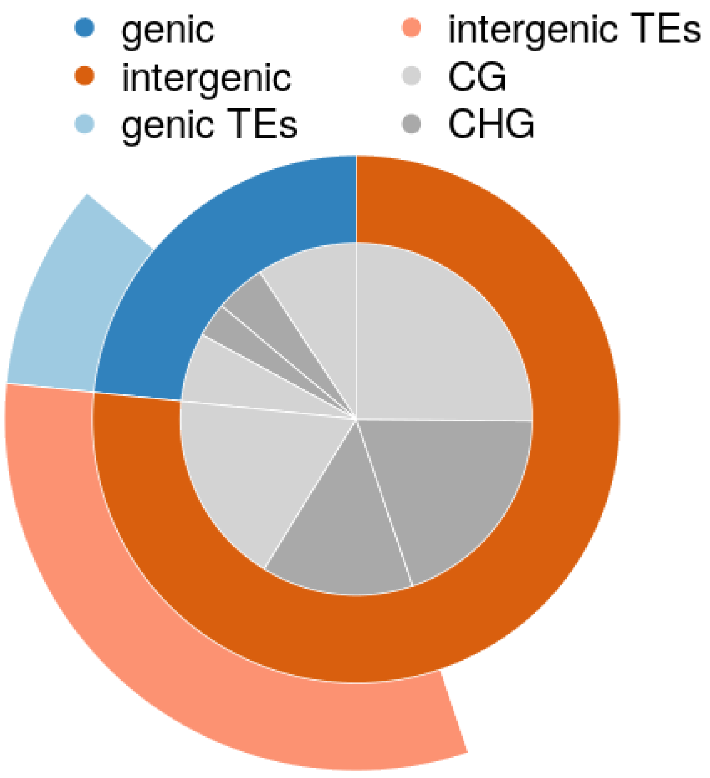


Figure S2

### Supplementary Figure S3

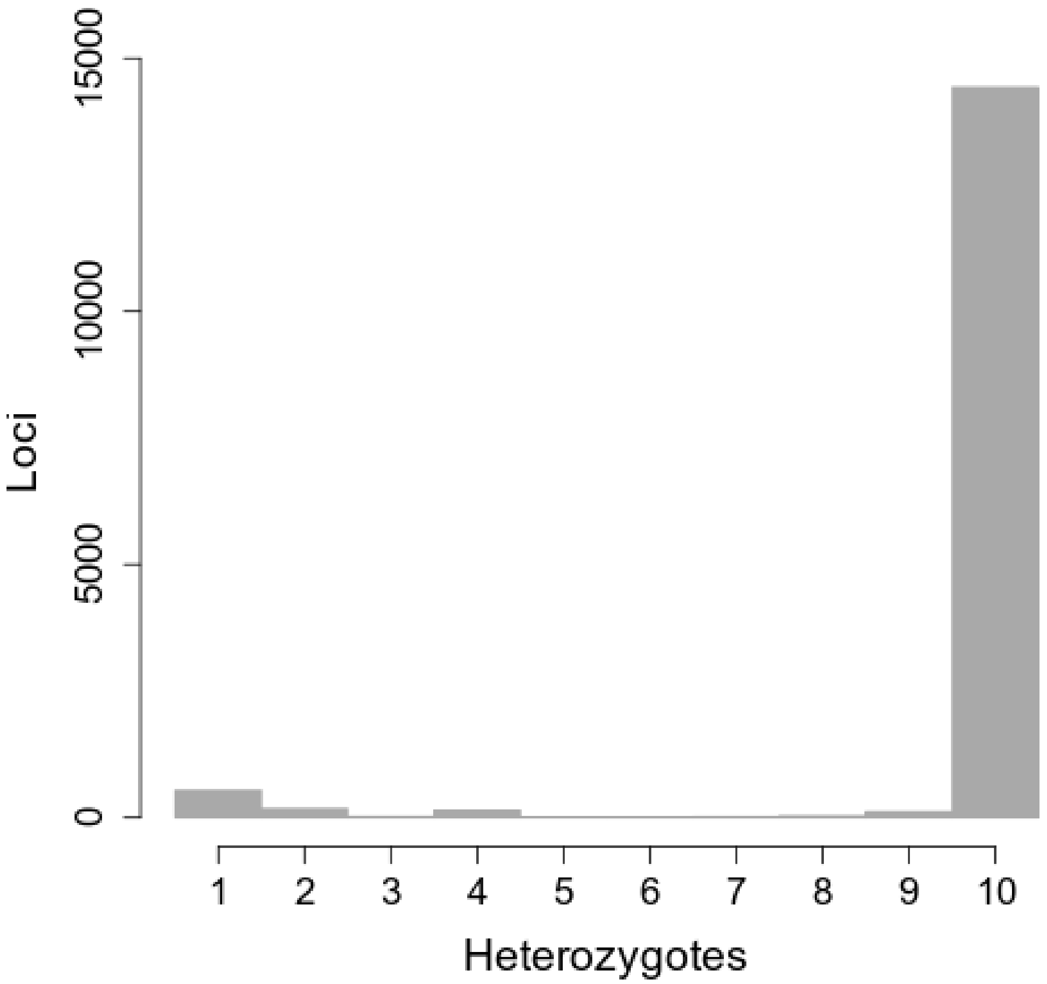


Figure S3

### Supplementary Figure S4

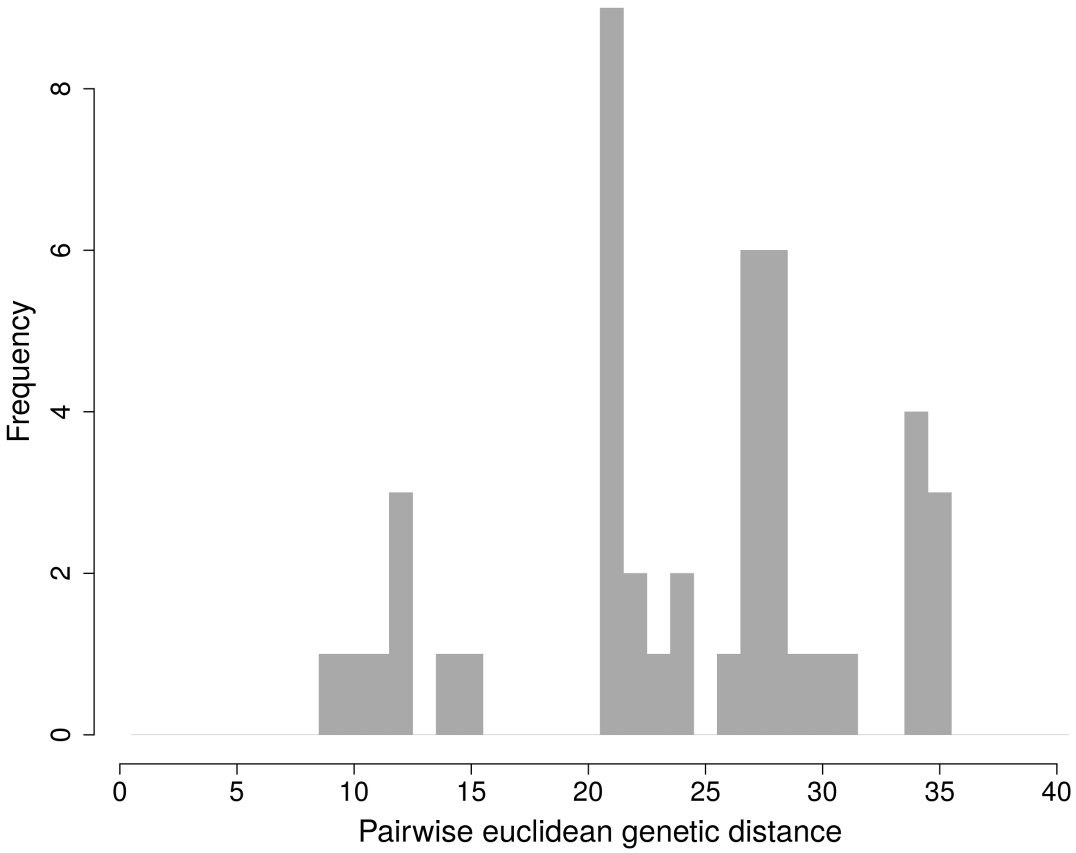

### Supplementary Figure S5

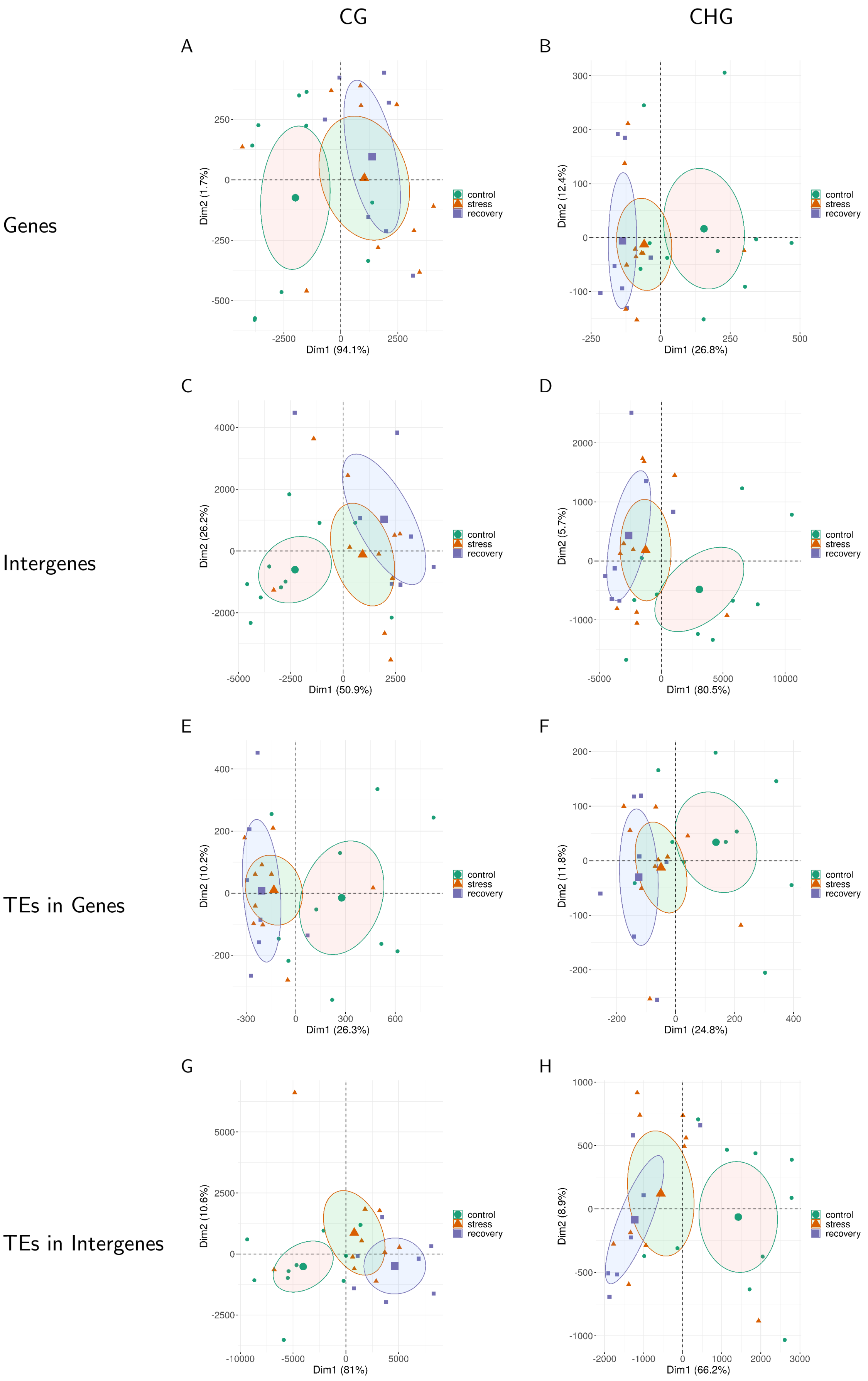
